# The impact of probiotic supplementation with effective microorganisms on water and gut microbiome of the *Common carp*

**DOI:** 10.1101/2024.12.14.628491

**Authors:** Michalina Jakimowicz, Tomasz Suchocki, Magda Mielczarek, Łukasz Napora-Rutkowski, Anita Brzoza, Teresa Kamińska-Gibas, Joanna Szyda

**Affiliations:** Wroclaw University of Environmental and Life Sciences, Department of Genetics, The Biostatistic Group, Kozuchowska 7, 51-631 Wroclaw, Poland; Polish Academy of Sciences in Golysz Institute of Ichthyobiology and Aquaculture Kalinowa 2, 43-520 Chybie, Poland

**Keywords:** Effective Microorganisms (EM), microbiome, 16S rRNA, gut, water

## Abstract

The efficacy of probiotic products in aquaculture covers multiple aspects that act directly on fish, such as improving resistance to fungal, bacterial, and viral infections, reproduction, feed efficiency, and overall growth performance, or indirectly through improved water quality. In our experiment, the application of two commercial water and two commercial food EM supplements was tested for their effect on water and the intestinal microbiome of common carp (*Cyprinus carpio*), as well as on carp growth performance. Probiotic supplementation did not result in a clear pattern of changes in overall microbial diversity or in the modification of the abundance of single genera. However, its effect was clearly manifested by an increase in the final weight of the fish. The most consistent alteration was the significant increase in Lactococcus abundance in the intestinal microbiome community compared to the control observed in both experimental setups. We conclude that the positive effect of *Lactococcus* on growth performance can be achieved by feed supplementation, which allows the increased abundance of Lactococcus in the fish intestine, which has a positive impact on glucose absorption and metabolism already demonstrated in the literature.

## 1. Introduction

Probiotics are living microorganisms (bacteria, yeast, and fungi) that, when provided in adequate amounts, provide health benefits to the host (Banerjee and Ray, 2017). The application of EM is extensive and extends from crop and animal husbandry to environmental protection, such as the deodorization of waste, the treatment of wastewater, and the control of eutrophication. In aquaculture, research has continuously increased with the demand for environmentally friendly aquaculture, focusing on improving water quality and alternatives to antibiotics (Gatesoupe, 1999). The efficacy of probiotic products in aquaculture covers multiple aspects that act directly on fish, such as improving resistance against fungal, bacterial, and viral infections, reproduction, feed efficiency, and overall growth performance, or through indirect impact through improving water quality (summarised e.g., by Banerjee and Ray, 2017a and El-Saadony et al., 2021a). Dawood and Koshio (2016) provided a review of the effects of probiotic supplementation specific to studies related to the *Carp* genus, demonstrating that research on the impact of probiotic supplementation in *Carp* breeding has been intensively carried out for almost two decades. Examples of recent studies using Common carp species as study material involve a positive effect of EM on growth reported by Jwher and Al-Sarhan (2022), observed increased activity of digestive enzymes and an improved immune response described by Mohammadian et al. (2022), a positive effect of *Lactobacillus acidophilus* on growth rate and immune response was demonstrated by Gabr et al. (2023) and of *Lactococcus* spp. by Feng et al. (2022), improved immune response due to supplementation with *Enterobacter asburiae* reported by Li et al. (2023), beneficial effects on the digestive system due to supplementation with *Lactobacillus rhamnosus* (Chen et al., 2024), as well as improved immunity, overall disease resistance and intestinal microbiome composition after supplementation of the fish diet with *Bacillus subtilis* (Xia et al., 2024).

Our research aimed to study the effect of different commercially available EM supplements on gut microbial communities of Common carp (*Cyprinus carpio*) and water, as well as on fish growth performance, by conducting a 94-day experiment. The microbiome composition was identified by sequencing two hypervariable regions (V3 and V4) of the gene that encodes the 16S rRNA ribosomal subunit.

## 2. Materials and Methods

### 2.1. The experiment

The experiment was conducted at the Institute of Ichthyobiology and Aquaculture in Golysz, Poland. It included seven 150-litre tanks from which water samples were obtained on day one and day 94 (last) of the experiment. The experimental setup comprised: (A1) - a control tank without fish and without probiotic supplementation, (A2, A3) - two water tanks with fish, without probiotic supplementation, (A4, A5) - two tanks with fish and the water supplement W1 and the feed supplement S1, (A6, A7) - two water tanks with fish and the water supplement W2 and the feed supplement S2. The EM supplements of water and food represented products commercially available in Poland, which according to the manufacturer’s information, contain seven species of *Lactobacillus* sp., three species of *Bifidobacterium* sp., *Carnobacterium divergens, Streptococcus salivarius*, and *Saccharomyces cerevisiae* (W1); all the above species plus *Lactobacillus delbrueckii, Bacillus subtilis, Rhodopseudomonas palustris*, and *Rhodobacter sphaeroides* (S1); *Lactobacillus plantarum, Lactobacillus casei*, and *Saccharomyces cerevisiae* plus species covered by manufacturer confidentiality (W2, S2). The experimental setup was visualised in Fig. 1. For all tanks, a constant water temperature of 25°C (+/-1°C) was maintained and every 7 days 15% of the water was refilled. Each tank A2-A7 was populated with five six-month-old individuals of common carp (*Cyprinus carpio*) with weights ranging from 16 g to 39 g, originating from the same mass spawning. On the last day of the experiment, three individuals were randomly selected from each tank, euthanised, and the contents of a small intestine was extracted from each of these individuals for sequencing.

**Fig. 1.**
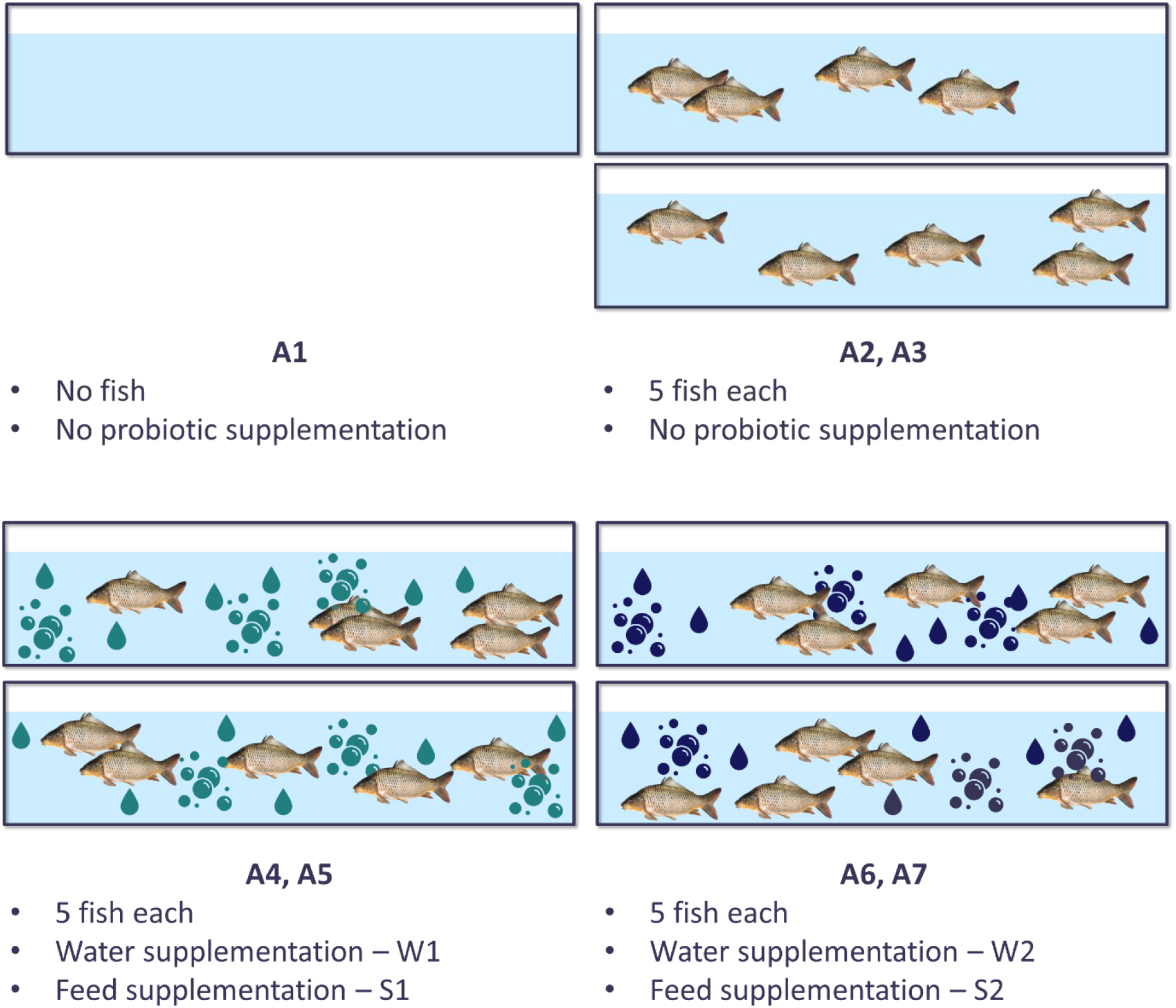
The experimental setup.

### 2.2. DNA handling and sequencing

DNA isolation from water (tanks A1-A7, two samples per tank, 14 samples) was carried out with the DNeasy PowerWater Kit (Qiagen). DNA isolation from small intestine content (tanks A2-A7, three fish per tank, 18 samples) was carried out with the DNeasy PowerFecal Pro DNA Kit (Qiagen). All samples were subjected to sequencing of the hypervariable regions V3 and V4 of the 16S rRNA gene with the MiSeq platform (Illumina) in the paired-end read mode with a read length of 300 bp. The regions were extracted and amplified using Bact 341F (5’ - CCT ACG GGN GGC WGC AG - 3’) and Bact 805R (5’ - GAC TAC HVG GGT ATC TAA TCC - 3’) primers. The library was created using the Herculase II Fusion DNA Polymerase Nextera XT Index V2 Kit with the 16S Metagenomic Sequencing Library Preparation Part #15044223 Rev. B protocol. Bioinformatic analysis of the amplicon sequences was performed using OmicsBox software version 2.2.4. First, the sequence quality was assessed using FastQC software version 0.11.8 (Andrews, 2010). Trimmomatic software version 0.38 (Bolger et al., 2014) was used to remove adapter sequences (TruSeq3) and trim all reads from their 3’ end to 270 bp length (i.e. by 30 bp). Furthermore, further read trimming was done by scanning each read with a four-base sliding window and trimming it when the average of the four-base qualities dropped below 20. Finally, reads with an average quality below 30 and shorter than 200 bp were discarded from downstream processing. The taxonomic classification of the remaining sequences was performed using Kraken Version 2.1.2 (Wood et al., 2019) with the confidence filter set to 0.05 and minimum hit groups of two using the combined Kraken and RefSeq databases implemented in the OmicsBox software. The Supplementary Table 1 presents the summary of sequence data for each environment.

### 2.3. Modelling the diversity of genera

Pairwise differences in the abundance of genera in water between the beginning and end of the experiment were tested with the partwise t-test and subjected to the Bonferroni correction (Dunnett, 1955). The diversity of genera within each sample (alpha diversity) was quantified with the Shannon index (Shannon and Weaver, 1949): 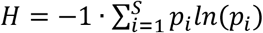, where *S* represents the number of considered genera and *p*_*i*_ is the frequency of the *i*-th genus in the sample. To test the significance of pairwise differences between alpha diversities, the Hutcheson t-test (Hutcheson, 1970) was used, for which the test statistics is defined as: 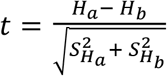, where *H*_*j*_ represents Shannon diversity index of the *j*-th sample and 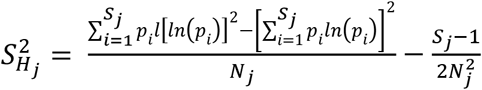 is the variance of *H*_*j*_ with *N*_*j*_ representing the number of reads in the *j*-th sample. Asymptotically, under the null hypothesis, the test follows the t distribution with the degrees of freedom equal to ^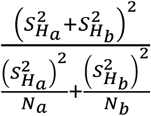^. The nominal P-values were corrected for multiple testing with Bonferroni correction. The differences in diversity between pairs of samples (beta diversity) were expressed as the Bray-Curtis distance (Bray and Curtis, 1957) 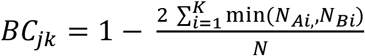, where *K* is the number of genera common between samples *A* and *B, N*_*Xi*_ is the number of reads representing the *i*-th genus in the sample *Xϵ*{*A, B*}, *N* represents the total number of reads over all genera in both samples. Both diversity measures were calculated using the vegan 2.6.8 R package (cran.r-project.org/web/packages/vegan/index.html), while the Hutcheson test statistic and corresponding nominal P-values were calculated using the ecolTest 0.0.1 R package (cran.r-project.org/web/packages/ecolTest/index.html).

In the second step, differences in abundance on a single genus level were estimated and tested using the Deseq2 package (Love et al., 2014). For each genus, log_2_FoldChange (LFC) was used to quantify the difference in abundance between samples, while *H*_0_ of no difference in abundance was tested using the Wald test 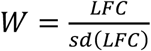 which asymptotically follows the standard normal distribution. The resulting nominal P-values corresponding to each tested genus were then expressed as False Discovery Rates (FDR, Benjamini and Hochberg, 1995) to account for multiple testing.

### 2.4. Modelling fish growth

One-way analysis of variance, fitting a tank as the experimental unit, was applied for testing *H*_0_ of no difference in fish weight between tanks. The first model was fitted to weights recorded on the first day of the experiment, and the second model to weights on the last (94) day of the experiment. Both models were fitted using the aov R function, followed by hypothesis testing based on the F test and the Duncan post hoc test implemented using the PostHocTest R function.

## 3. Results

### 3.1. Most abundant genera

The analysis was carried out on the genus level. The composition of the predominant genera in the water at the beginning of the experiment was similar, although not identical, between tanks. Only *Devosia* (from 0.17% to 36.20%) and *Nordella* (from 1.06% to 21.80%) were among the top four abundant genera in the six tanks. *Aquirufa* was among the four most abundant species in both control tanks A2, A3 and experimental tanks A4, A5 with a relative abundance varying between 0.23% and 32.60%, while *Cetobacterium* was highly abundant in control tanks A2, A3 and experimental tanks A6, A7 (Fig. 2). At the end of the experiment, the set of predominant genera was different from that at the beginning. *Cetobacterium* (3.54% to 22.00%) and *Nocardidides* (3.44% to 12.20%) were among the four most abundant in all tanks. *Rhodanobacter* was among the most abundant genera in all experimental tanks A4, A5, A6, and A7, *Candidatus Koribacter* was highly abundant both in the control A2 (46.00%), A3 (45.00%) as well as in one experimental tank A4 (30.00%), *Singulsiphera* was highly abundant only in W2S2 (from 6.32% and 6.81%) (Fig. 3). Additionally, no significant pairwise differences in abundance of the top 100 genera were detected between the end and the beginning of the experiment.

**Fig. 2.**
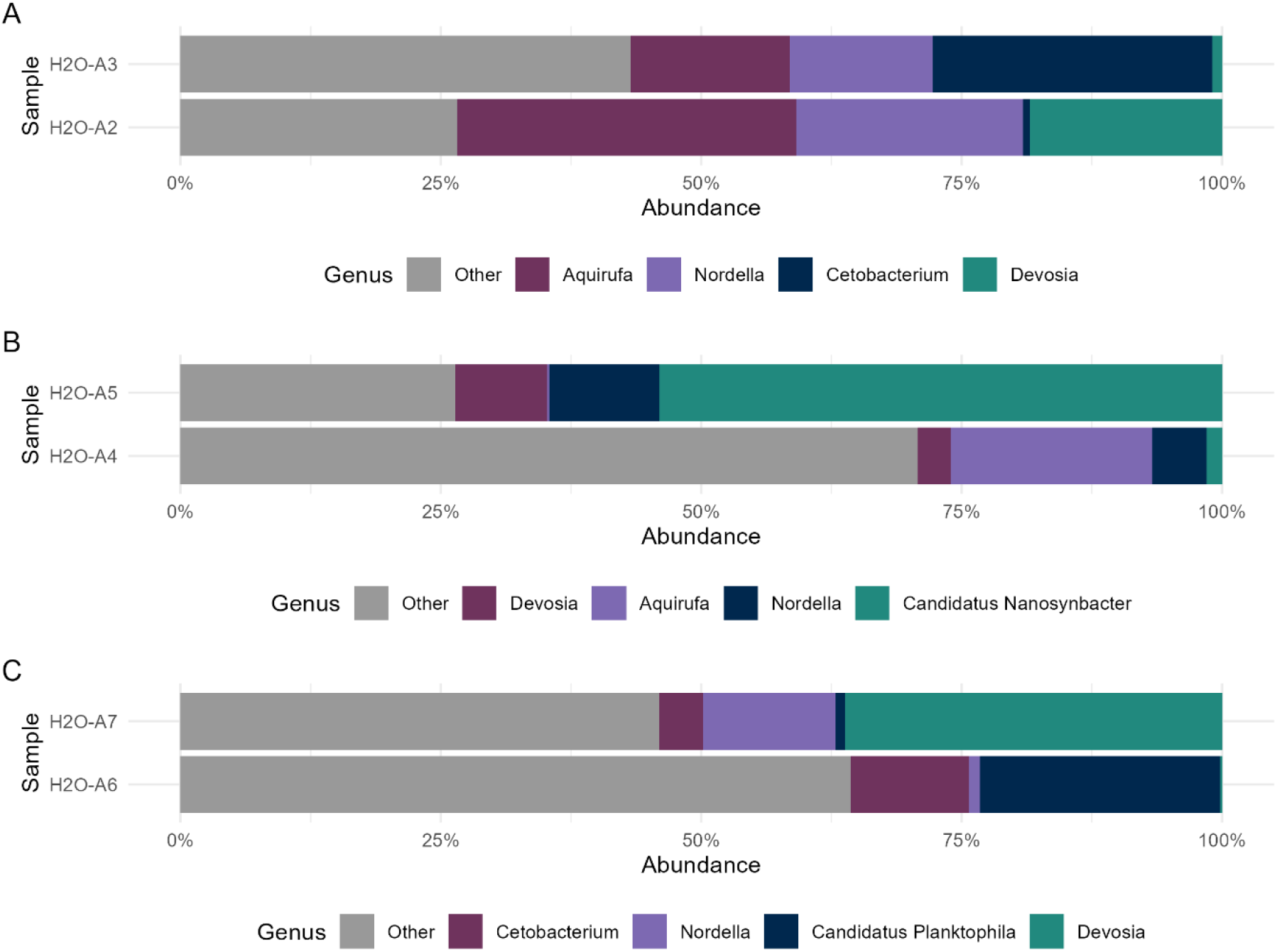
Relative abundance of genera in water samples at the beginning of the experiment. A - control group, B - W1S1, C - W2S2.

**Fig. 3.**
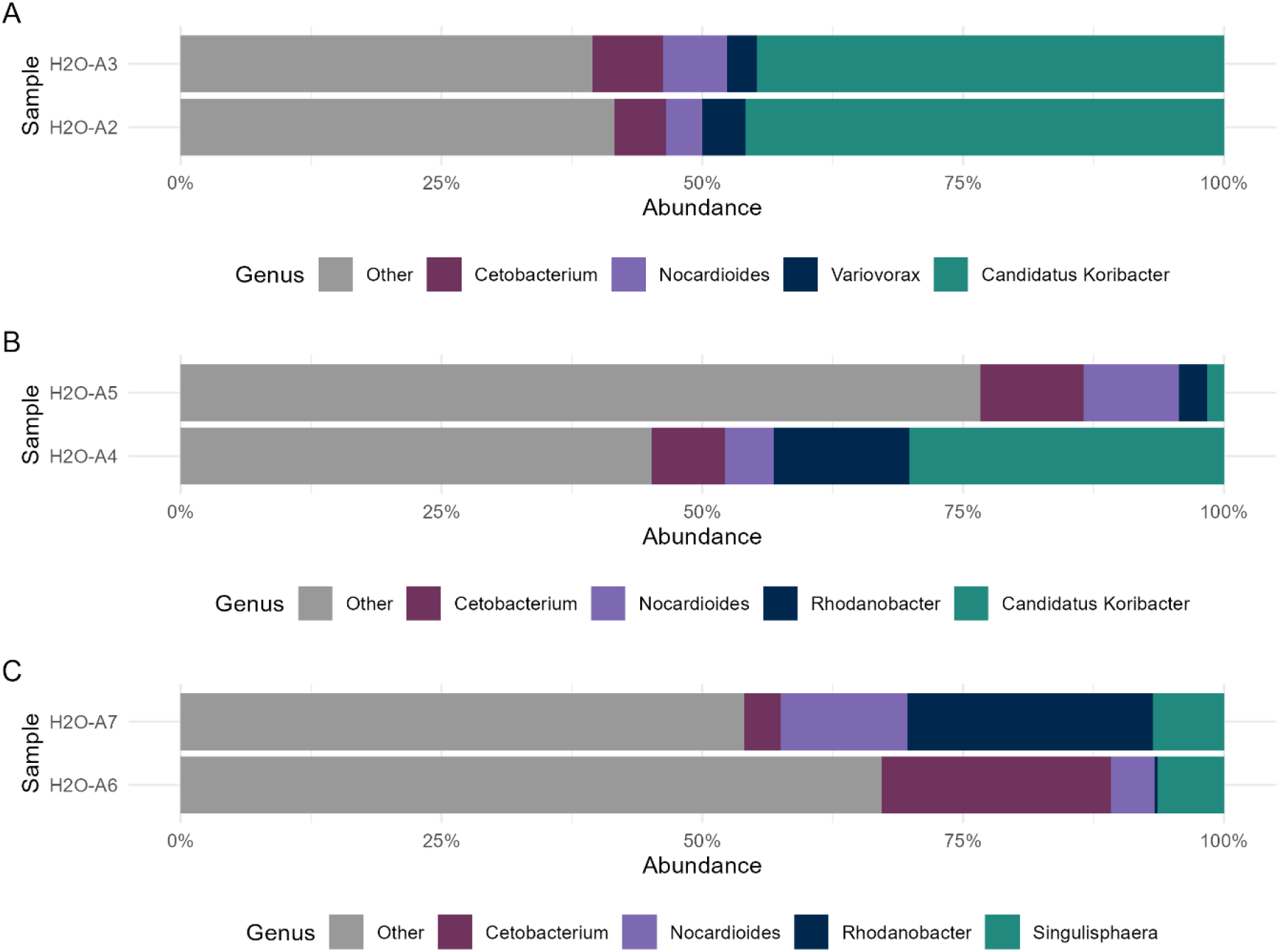
Relative abundance of genera in water samples at the end of the experiment. A control group, B - W1S1, C - W2S2.

Regarding the microbial composition of the intestine, the four genera that prevailed the most in all samples were *Acinetobacter* (from 0.02% to 27.50%), *Cetobacterium* (from 1.51% to 78.70%), *Lactobacillus* (from 0.68% to 25.60%), and *Latilactobacillus* (from 0.16% to 16.50%). Still, as visualised in Fig. 4, there is a considerable variation in the abundance of the most common genera, so even those four generally most abundant genera were not present among the top four in each sample. However, no tank- or experimental-design-related pattern emerges.

**Fig. 4.**
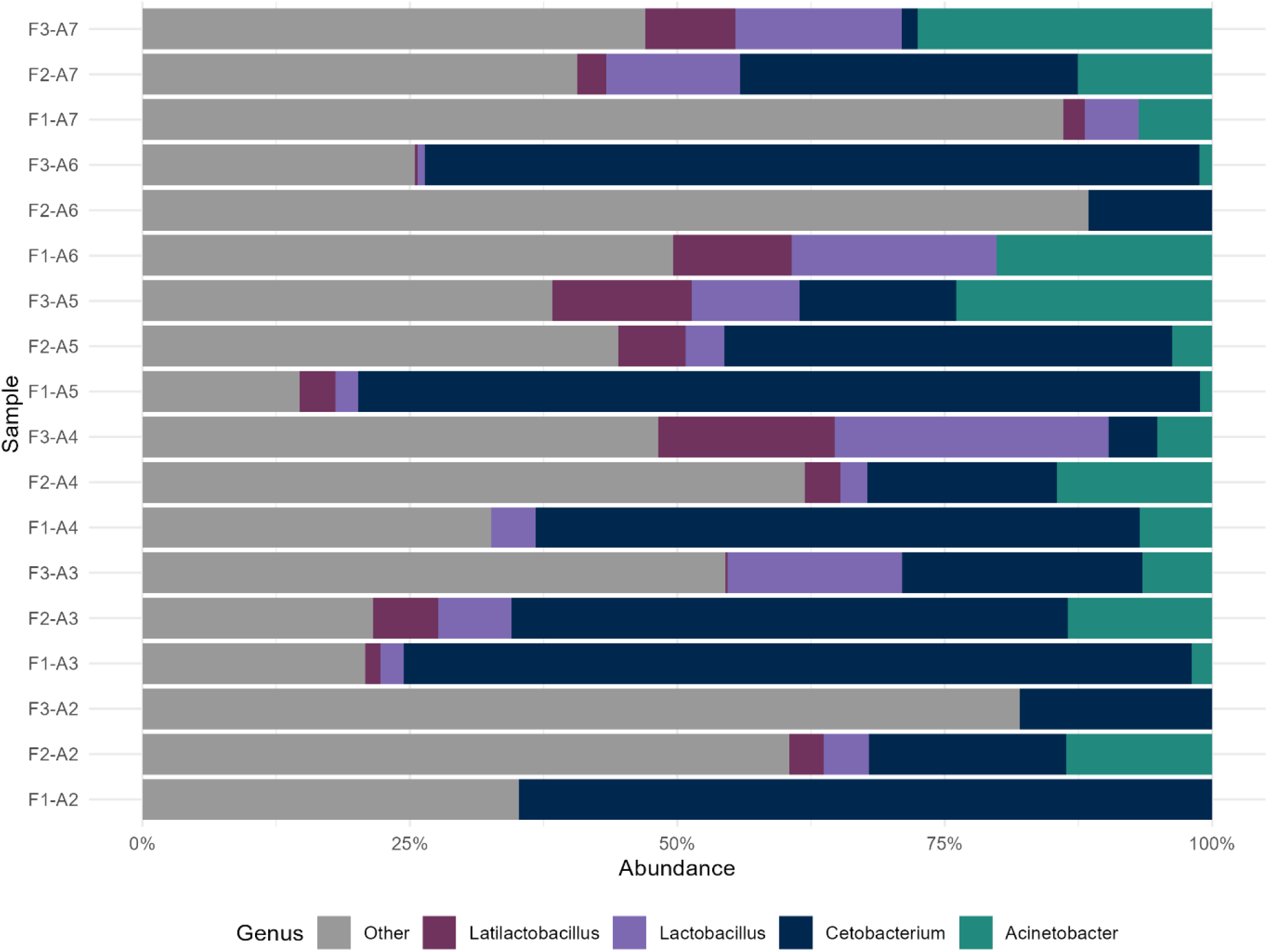
Relative abundance of genera in intestinal samples at the end of the experiment. F represents fish, A represents tank.

### 3.2. Diversity of genera

The diversity of genera within the water samples expressed by the Shannon index varied considerably between the tanks Fig. 5 as did the diversities within the intestinal samples (Fig. 6). In general, the diversity within the water samples that ranged between 2.71 for A3 and 3.67 for A5 with a mean of 3.09 was higher than within the intestinal samples that varied between 1.07 (F1-A5) and 2.86 (F3-A3), with a mean of 2.21. However, almost all pairwise comparisons revealed significant differences in alpha diversities between samples. Supplementation did not result in a consistent pattern of increase or decrease of diversity, in the sense that for a control sample A3 and a W1S1 sample A4 a lower alpha diversity was observed at the end, for other samples – at the beginning phase of the experiment. The highest increase in variability was observed in the experimental sample A5 for which the Shannon index increased by 1.65 (P=0.00011). Similarly, for intestinal samples only four pairwise comparisons of within sample diversities resulted in a significant difference and involved individuals from A3-A5, A3-A6, A5-A6 and within A6, but no characteristic pattern emerged. Within-sample diversity estimates were summarized in Supplementary Table 1 and the results of all pairwise comparisons in Supplementary Table 2.

**Fig. 5.**
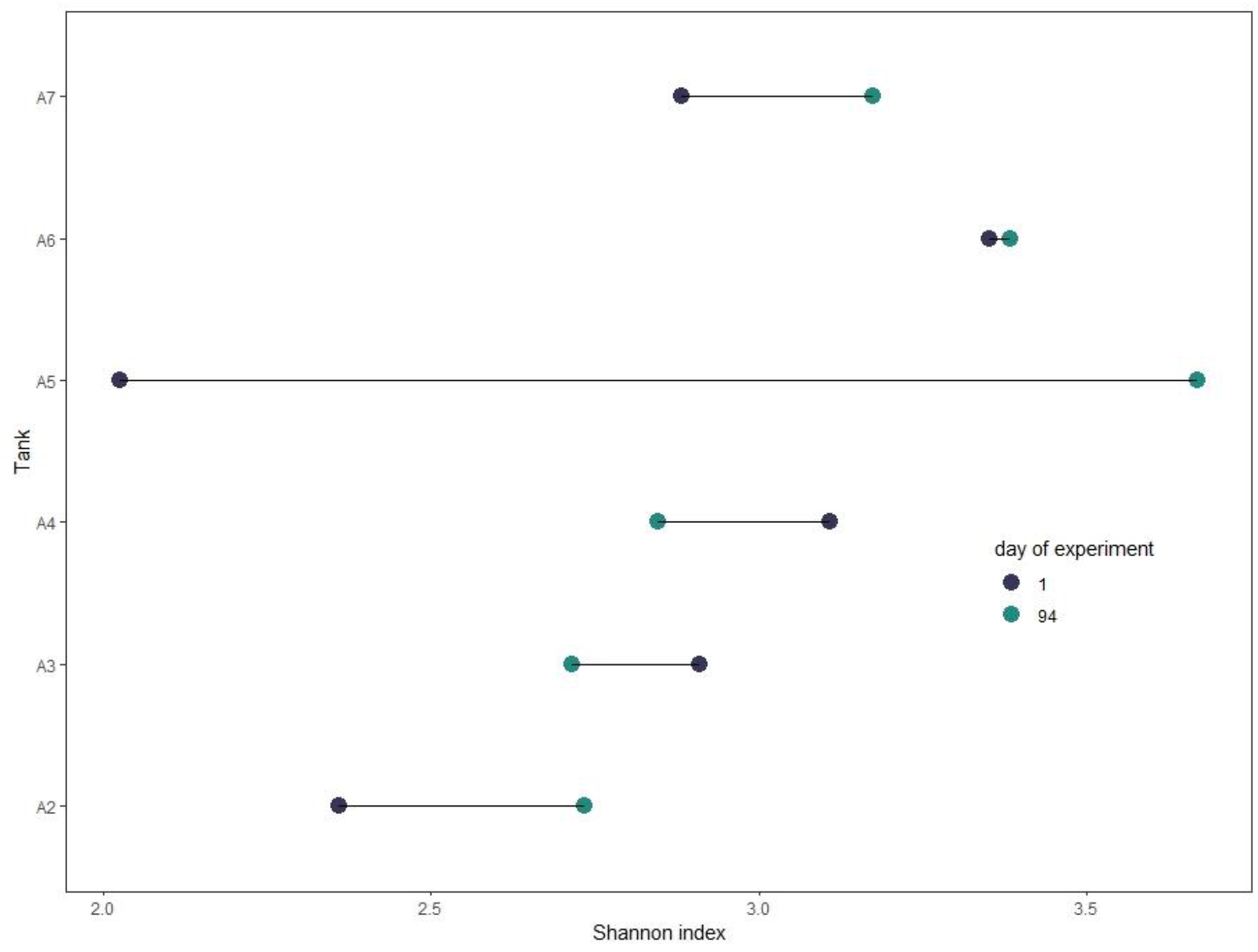
Diversity within water samples quantified by Shannon index at the beginning and end of the experiment.

**Fig. 6.**
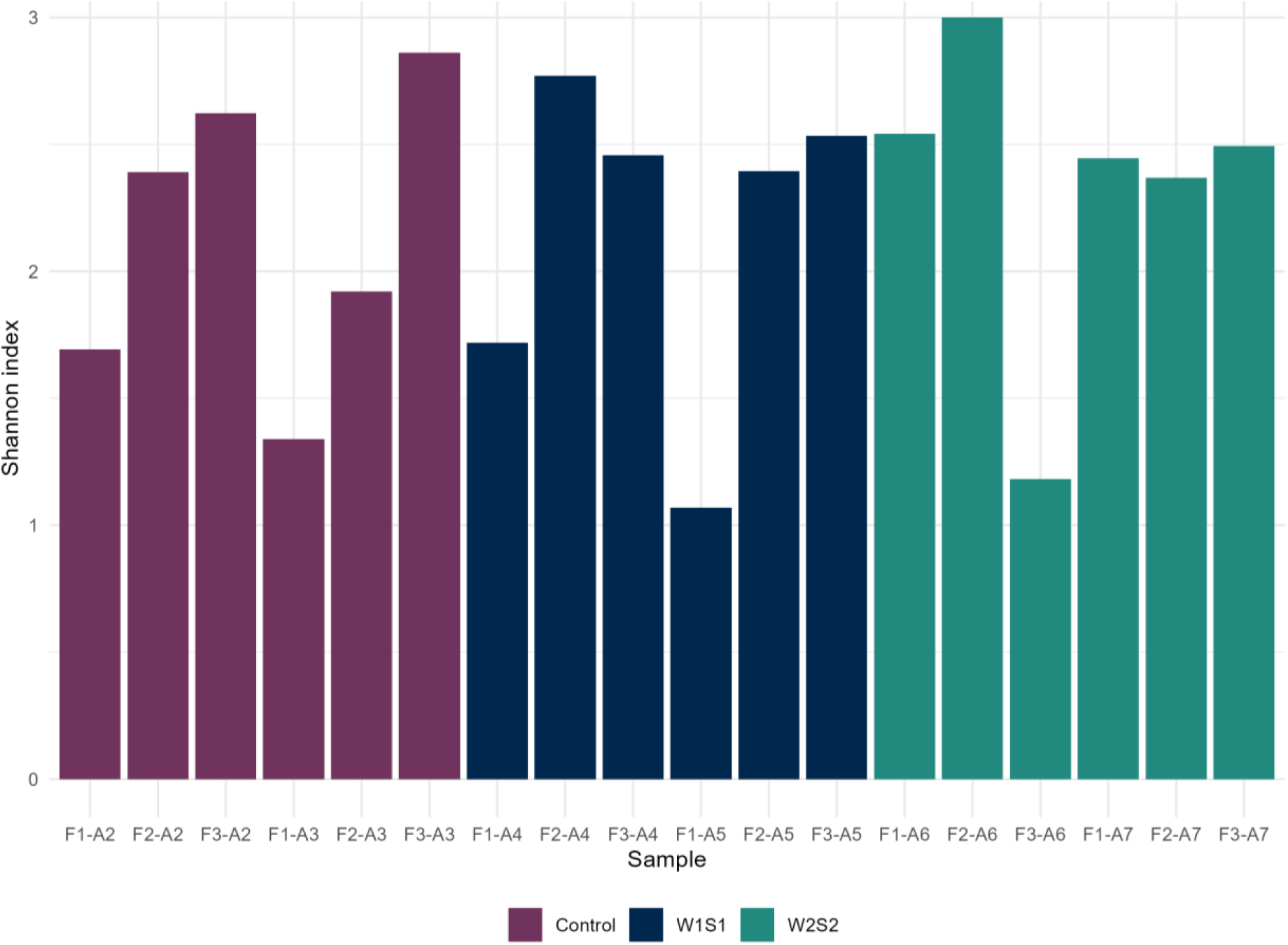
Diversity within intestinal samples quantified by Shannon index. F - fish, A - tank.

The pairwise differences between the abundance of genera in water samples on the last day of the experiment, expressed by the Bray – Curtis pairwise distances visualized in Fig. 7, were generally high, with the mean distance equal to 0.62 and for all but one pairwise sample comparison, exceeded 0.50. However, no systematic pattern of between sample diversities emerges. The most divergent water samples (BC=0.82) were A3 and A4, representing the control and the experimental setup W2S2, while both control samples, A2 and A3, were the most similar (BC=0.42). Also, there was no clear pattern of pairwise diversities between intestinal samples (Fig. 8). The longest distance of 0.98 was estimated between an individual sample from the control group A2 and an individual from the experimental setup W2S2, which also reflected the highest diversity between the water microbial composition of those samples. Surprisingly, the most similar microbial composition expressed by BC of 0.24 was estimated between samples of fish representing different experimental setups, W1S1 and W2S2, kept in tanks A5 and A6.

**Fig. 7.**
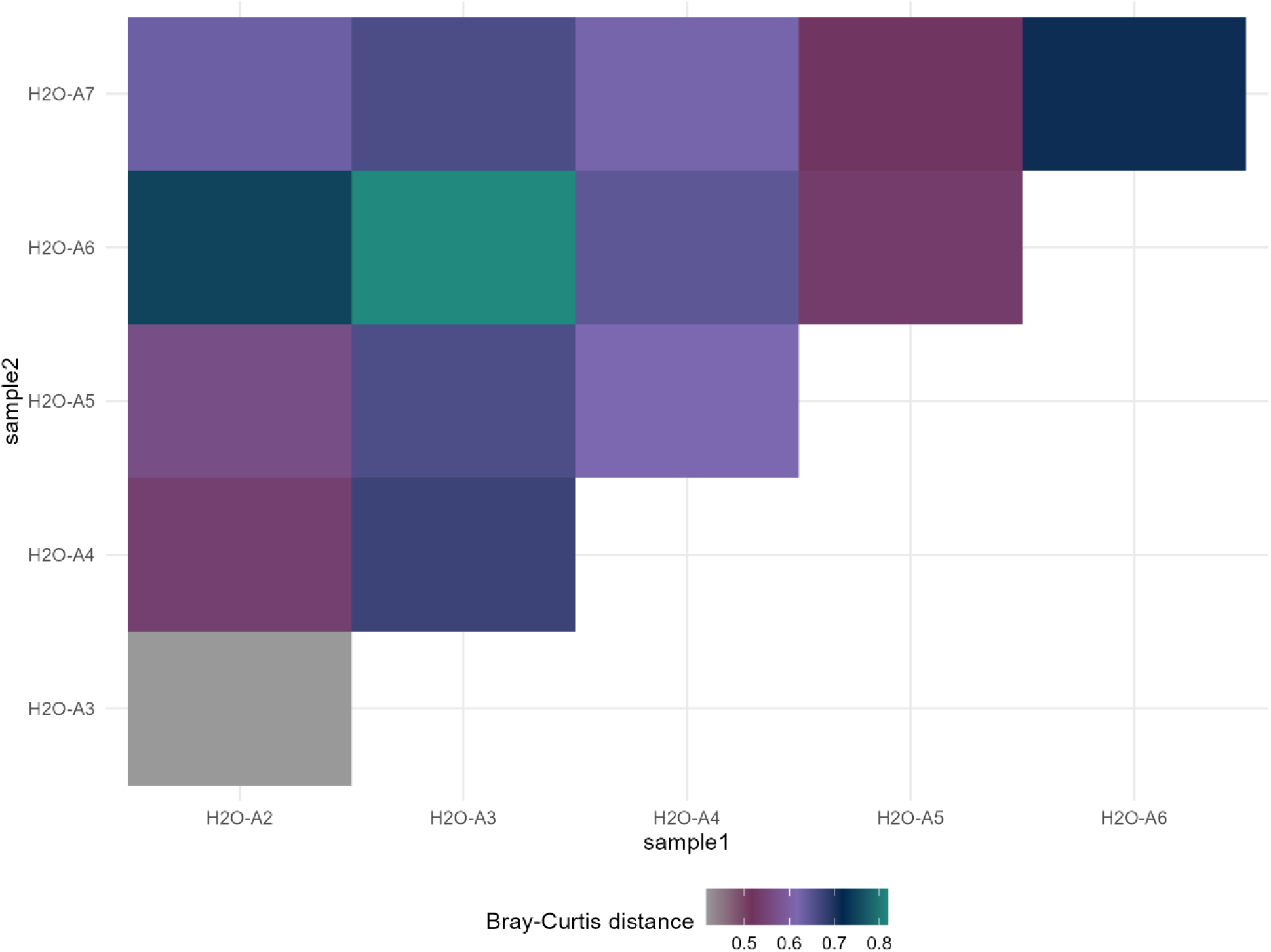
Pairwise distances between water sample genera abundances at the end of the experiment expressed by the Bray – Curtis metric.

**Fig. 8.**
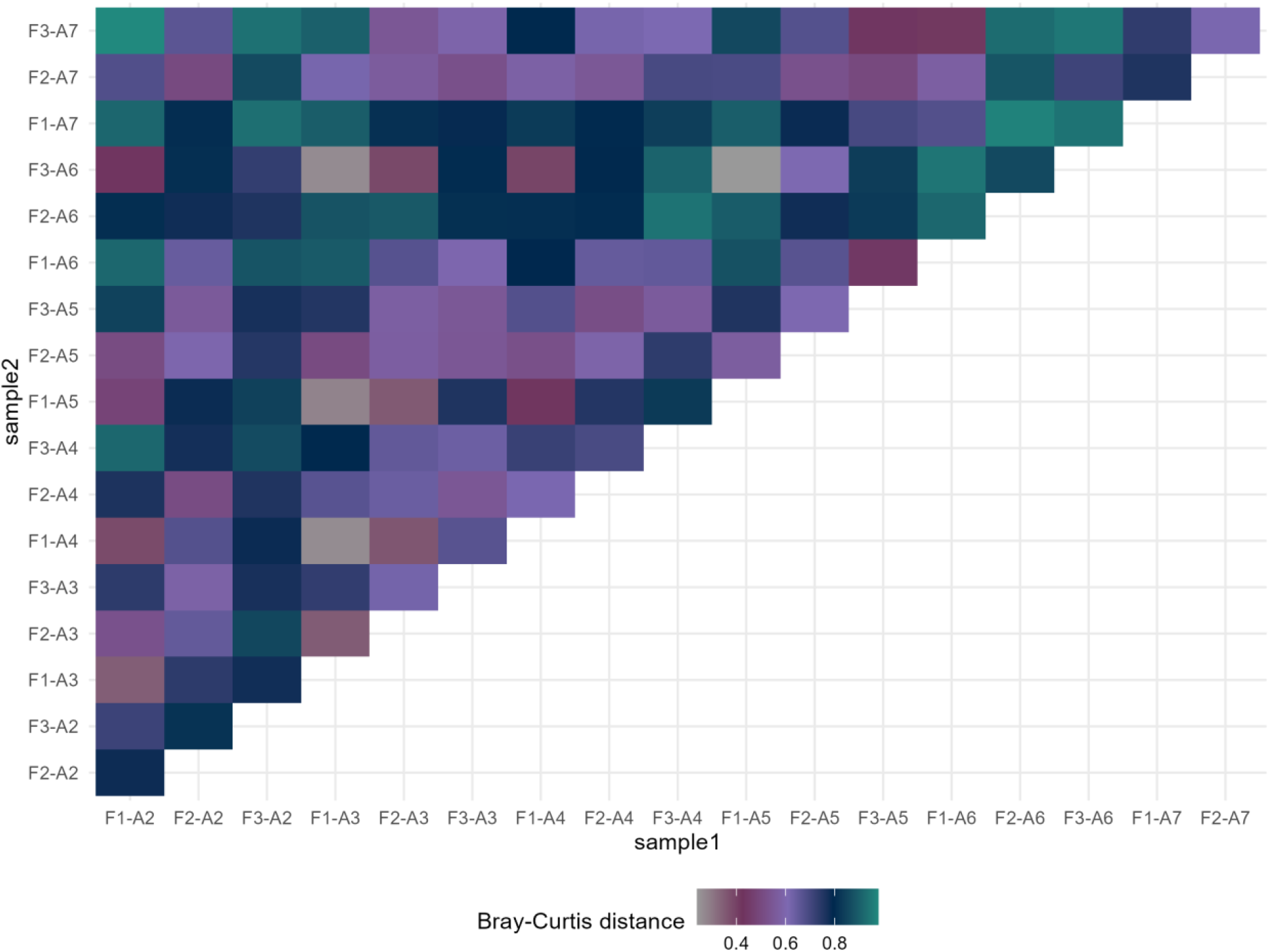
Pairwise distances between intestinal sample genera abundances expressed by the Bray – Curtis metric. F represents fish, A represents the experimental design.

**Fig. 9.**
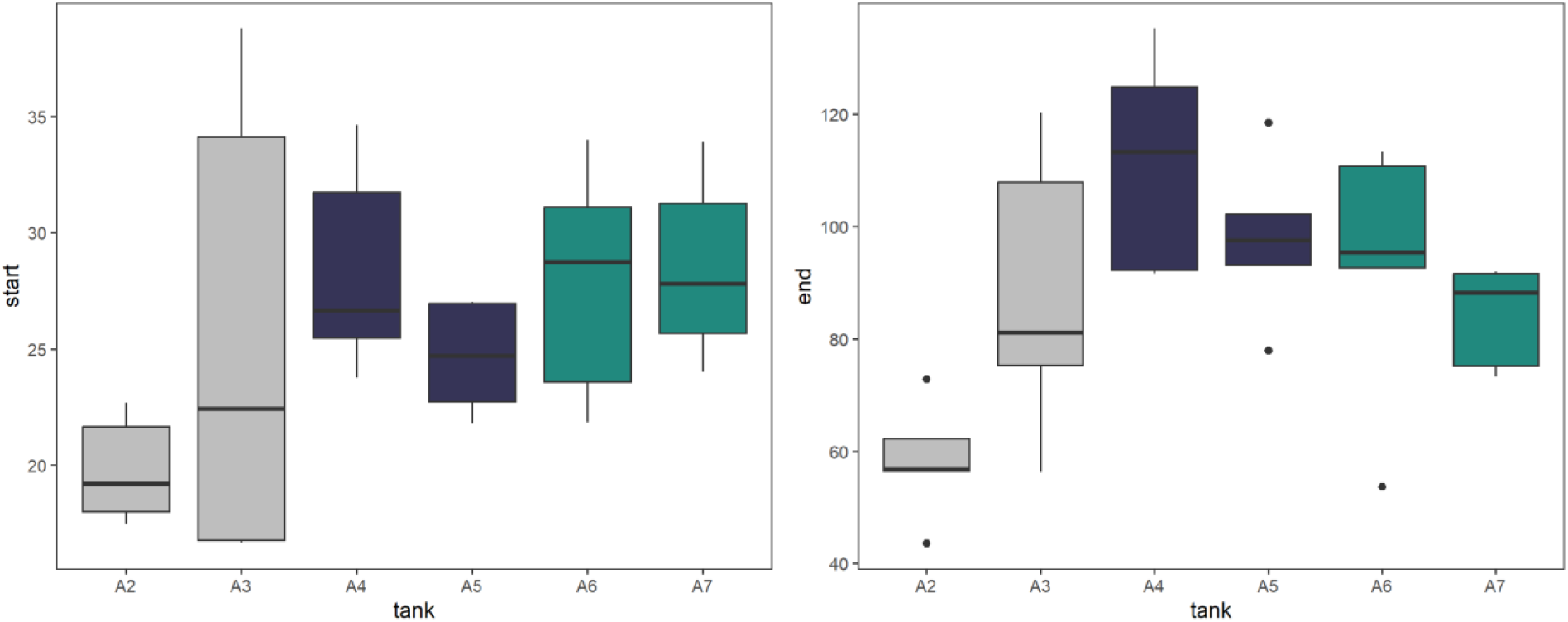
Fish weights at the start (day 1) and end (day 94) of the experiment.

### 3.3. Differences in abundance of single genera

Since no clear pattern emerged for overall microbial diversity, the second part of the study focused on particular genera.

The differential abundance test revealed that despite the unified conditions of the experimental setup, there were still significant differences in the microbial compositions of the tank water, especially between the control and W1S1 tanks: *Candidatus Planktophila, Vibrio, Candidatus Nanosynbacter, Emticicia, Polynucleobacter, Novosphingobium, Candidatus Nanopelagicus*, but also between the control and W2S2 tanks *Candidatus Planktophila and Candidatus Nanopelagicus*. At the end of the experiment, regardless of the supplementation, the microbial composition of the water was stabilised, so only *Polynucleobacter* remained significantly (FDR=0.049) more abundant in W1S1 compared to the control samples. In the intestinal microbiome, W1S1 supplementation led to a significant decrease in *Candidatus Koribacter* (FDR=9.70·10^-15^), *Planctopirus* (FDR=4.19·10^-12^), *Iamia* (FDR=5.41·10^-10^), and *Singulisphaera* (FDR=7.94·10^-13^). The significant effects of W2S2 supplementation were more complex, in a way that the abundances of *Leptolyngbya* (FDR=1.73·10^-12^) and *Ruminococcus* (FDR=1.21·10^-13^) decreased after supplementation, while the abundances of *Aquicella* (FDR=4.90·10^-13^) and *Intestinimonas* (FDR=8.64·10^-13^) significantly increased. Only the abundance of *Lactococcus* (FDR for W1S1 =3.10·10^-16^ and FDR for W2S2 =4.57·10^-21^) significantly increased regardless of the supplementation setup.

### 3.4. Fish weight

Fish weights at the beginning and end of the experiment were visualized in Figure 8. On day 1, no significant differences in weight between tanks were observed (P=0.130), but after 94 days, significant differences between tanks emerged (P=0.004). In particular, weights significantly higher than in the control tank A2 were recorded for the experimental tanks A4 (P=0.002) and A5 (P=0.025), both subjected to W1S1 supplementation.

## 4. Discussion

In the literature, there is general agreement that the application of probiotic water supplementation with commercial products does not influence the microbiome composition of water (Tang et al., 2016; Zhou et al., 2017; Zheng et al., 2017). The authors argued that this is due to problems with the successful colonization of the supplemented bacteria in the established, natural macrobiotic water community. Due to the lack of detailed information on the composition of the products implemented in our study, this hypothesis could not be formally tested. However, this situation can indirectly be concluded from the absence of significant differences (except for *Polynucleobacter*) between the control and experimental tanks on the last day of the experiment, which was also observed in our study. Furthermore, the significant variation between tanks in the abundance of genera at the beginning of the experiment indicates that despite the unified conditions, time is needed for the microbial community to stabilize after colonization of the tanks at the beginning of the experiment.

Tang et al. (2016) and Zheng et al. (2017) did not observe a significant effect of water supplementation on growth performance. In our experiment, it was evident that, as discussed above. However, the water microbial community remained unchanged, the supplementation significantly affected the intestinal microbiome of fish, which may have led to improved growth performance. This implies that growth performance is not affected by probiotic water supplementation, but by directly supplementing feed. In particular, our study demonstrated a significant increase in the abundance of *Lactococcus* in intestinal microbiome communities of fish kept in tanks subjected to probiotic supplementation. A positive effect of *Lactococcus* on the growth performance of *Common carp* (and other aquaculture species) was previously reported by Feng et al. (2019, 2022) and Zhang et al. (2019), who suggested that the effect is due to a positive influence of the genus on glucose absorption and metabolism.

## 5. Conclusion

Although probiotic supplementation did not cause any significant modifications in the water or intestinal microbiome of the fish, the effect of supplementation was clearly manifested by an increase in the final weight of the fish. Based on the literature that, on the one hand, has widely reported no effect of water supplementation on the microbial composition of water and, on the other hand, has demonstrated a positive impact of *Lactococcus* on fish growth performance, we conclude that the positive effect on growth performance can be achieved by feed supplementation which allows the enhancement of the abundance of *Lactococcus* in the fish intestine. Furthermore, following the hypothesis raised by Moroni et al. (2021), *Lactococcus* itself may also exhibit a positive effect on the composition of the fish intestinal microbiome that is secondary and not directly due to food supplementation.

## Acknowledgements

Calculations were carried out at the Wroclaw Centre for Networking and Supercomputing and Poznan Supercomputing and Networking Center.

